# Comparing efficiencies of population control methods for responding to foreign animal disease threats in wild pigs

**DOI:** 10.1101/2024.07.26.605354

**Authors:** Nathan P. Snow, Benjamin Smith, Michael J. Lavelle, Michael P. Glow, Kayleigh Chalkowski, Bruce R. Leland, Sarah Sherburne, Justin W. Fischer, Keely J. Kohen, Seth M. Cook, Hatton Smith, Kurt C. VerCauteren, Ryan S. Miller, Kim M. Pepin

**Affiliations:** USDA/APHIS/ Wildlife Services, National Wildlife Research Center, 4101 LaPorte Ave., Fort Collins, Colorado 80521, USA; USDA/APHIS/ Wildlife Services, 5730 Northwest Pkwy #700, San Antonio, Texas 78249, USA; USDA/APHIS/ Veterinary Services, Center for Epidemiology and Animal Health, 2150 Centre Avenue, Fort Collins, Colorado, 80526, USA

**Keywords:** Aerial, African swine fever, Eradication, Feral swine, Removal, *Sus scrofa*, Trapping, Toxicant, Ungulates, Wild boar

## Abstract

Introductions of foreign animal diseases (FADs) into free-ranging wildlife can be difficult to control and devastating for domestic livestock trade. Combating a new FAD introduction in wildlife with an emergency response requires quickly limiting spread of the disease by intensely removing wild animals and recovering their carcasses for proper disposal. In the case of African swine fever virus (ASFv) in wild pigs (*Sus scrofa*), which has been spreading in many regions of the world, there is little information on the time- and cost-efficiency of methods for intensively and consistently removing wild pigs and recovering carcasses in an emergency response scenario. We compared the efficiencies of aerial operations, trapping, an experimental toxic bait, and ground shooting in northcentral Texas, USA during two months in 2023. Removing and recovering carcasses of wild pigs averaged a rate of 0.15 wild pigs/person hour and cost an average of $233.04/wild pig ($USD 2023) across all four methods. Aerial operations required the greatest initial investment but subsequently was the most time- and cost-efficient, costing an average of $7,266 to incrementally reduce the population by 10% including recovering carcasses. Aerial operations required a ground crew of ∼7 people/helicopter to recover carcasses. Costs for reducing the population of wild pigs using trapping were similar, although took 13.5 times longer to accomplish. A benefit of trapping was carcass recovery was incorporated. Toxic baiting was less efficient because carcass recovery required substantial time, and we removed very few wild pigs with ground shooting in this landscape. We recommend combining aerial and trapping methodologies to remove wild pigs and their carcasses efficiently and effectively during a FAD response. Overall, our findings can inform the preparation of resources, personnel needs, and deployment readiness for FAD responses involving wild pigs.

## Introduction

Foreign animal disease (FAD) introductions have occurred at increasing rates in recent years because of faster and cheaper global trade and transportation (Baker et al., 2022). Rapid and intensive response efforts to eradicate newly introduced diseases early are critical (Miller et al., 2022; Pepin et al., 2022) to avoid economic, public, and animal health consequences (Paarlberg et al., 2008; Negesso et al., 2016; McElwain and Thumbi, 2017). Emergency response plans include strategies for eradicating FADs in wild animal populations especially when the disease can be vectored by wild animal hosts to domestic animals (Smith et al., 2007; Miller and Pepin, 2019; Danzetta et al., 2020). Efforts to control or eradicate FADs in wild animals are immense, including intensive reductions in free-ranging populations, removal of carcasses, exhaustive surveillance, development of widely distributable vaccines, and long stretches of fencing (Smith et al., 2007; Danzetta et al., 2020; Dressel et al., 2024).

Large-scale, rapid, and intensive removal of wild animals is difficult because it requires immediate mobilization of collaborative resources, often at a national level (Lefrançois et al., 2023). Successful implementation of such emergency responses requires *a priori* operational knowledge to better coordinate data collection to inform decision-making during the response (Lefrançois et al., 2023; Zinsstag et al., 2023). Among the key steps to planning an emergency response is allocating resources to achieve desired outcomes (FEMA, 2010). Despite the expense, intensive removal of infected wild populations can have a greater cost-benefit than no action during an FAD outbreak (Smith et al., 2007).

Wild ungulates represent significant challenges for FAD responses. Wild ungulates exist or have been introduced around the world (Lever, 1985; Nelle, 1992; Lever and Yalden, 1995; Jaksic et al., 2002; Long, 2003; Hulme et al., 2008; VerCauteren et al., 2019) and are reservoirs for a wide range of animal diseases worldwide that can be transmitted to domestic animals (Martin et al. 2011). Wild ungulates can be “tricky species” during emergency response due to a lack of effective tools to quickly reduce their populations, quick adaptation to avoid control efforts, are frequently neophobic, and regulatory limitations on available tools or approaches (Parkes and Panetta, 2009). A current example of a spreading FAD in wild ungulates that is important to food security and international trade is African Swine Fever virus (ASFv). This virus is a highly contagious and deadly disease for domestic swine (*S. scrofa domesticus*), European wild boar (*S. scrofa*), and wild pigs (*S. scrofa*) and is spreading across Europe, Asia, and the Caribbean (Penrith and Vosloo, 2009; Cwynar et al., 2019; Sur, 2019; Gonzales et al., 2021). In countries that are currently ASFv-free (e.g., USA, Canada, Australia, and others), there have been ongoing efforts to prevent and respond to potential introductions of ASFv into wild pig populations (e.g., Pepin et al., 2020; Bradhurst et al., 2021; MacDonald and Brook, 2023; Brown et al., 2024). If introduced into wild pigs, intensive reduction of wild pig populations is recommended given they represent a high-risk pathway for endemic persistence (Brown et al., 2021). In addition, recovering and disposing of carcasses of wild pigs following control efforts would be necessary (Zani et al., 2020), which can be challenging depending on the control technique and environmental conditions. Identifying the most time- and cost-effective methods for rapidly reducing populations of wild pigs and recovering carcasses can provide valuable planning information for preventing establishment of ASFv.

No single methodology can reduce populations of wild pigs in all situations (Ditchkoff and Bodenchuk, 2020). Four of the most effective and widely deployable removal methods are aerial shooting, trapping, toxic baiting, and ground shooting. Aerial shooting from a helicopter or fixed-wing aircraft can quickly cover large tracts of land and has been shown to be highly time-effective (Bodenchuk, 2014; Davis et al., 2018), but may pose challenges in situations where carcasses also need to be recovered and removed. Advances in trap designs (e.g., cellular traps; Ditchkoff and Bodenchuk, 2020; Gaskamp et al., 2021) and strategies (e.g., whole sounder removal; Lewis et al., 2022; Kilgo et al., 2023) have made substantial progress toward effectively reducing populations of wild pigs but require extensive labor and time. Toxic baits are not currently registered for use in some countries but are used in other countries with extensive non-native populations of wild pigs (e.g., Australia and New Zealand) and are under development in USA (Snow et al., 2017a; Poché et al., 2019). Toxic baits can be effective at removing large proportions of wild pigs with less labor than trapping (e.g., Snow et al., 2019b; Snow et al., 2021b), but have inherent risks to other species (Beasley et al., 2021; Snow et al., 2021a) and generate carcasses of wild pigs that are difficult to locate (Poché et al., 2018; Snow et al., 2021b). Ground shooting has been shown to be cost-effective under certain scenarios (Hamnett et al., 2023) and is frequently used to target individual animals, but utility is likely dependent on terrain and landcover.

In the context of an emergency response to a FAD introduction such as ASFv, multiple removal methods will likely be used but time- and cost-efficiency under intense-deployment remains poorly understood. Without a better understanding, it is difficult to proactively and adequately prepare response efforts including resource stockpiling, budget and personnel needs, and a realistic estimation of how rapidly the population of wild animals can be removed relative to ongoing transmission of disease. Our objective was to compare the time and costs required for the four common methods of removing wild pigs (i.e., aerial operations, trapping, toxic baiting, and ground shooting) and recovering carcasses during an intensive population control effort. Secondly, we compared the impact each of the removal methods had on the population density of wild pigs. Ultimately, our goal is to inform emergency response readiness for preventing establishment or eliminating FADs in a free-ranging population of wild pigs.

## Methods

### Study area

We conducted our study on a 225.1 km^2^ portion of an active cattle ranch located in north-central Texas, US during January–May 2023. The landscape is characterized as part of the southwestern tablelands of the south central semi-arid prairies ecoregion (Bailey, 1980), dominated by shortgrass and midgrass prairies, mesquite (*Prosopis* spp.) savanna, cedar (*Juniperus* spp.), riparian areas of plains cottonwood (*Populus* spp.), and interspersed wheat fields. The topography consists of broad, rolling plains with interspersed elevated tablelands, red-hued canyons, badlands, and dissecting river breaks. During our study, temperatures averaged 7–21 °C and 97.2 mm of precipitation occurred (www.ncdc.noaa.gov/cdo-web/search). Management of wild pigs prior to our study was minimal, with occasional hunting and trapping primarily for recreational purposes. The landscape was estimated to hold a potential density of 3–5 wild pigs/km^2^ (Lewis et al., 2019).

### Study design

We divided the study area into four zones (ranging from 46.1–64.2 km^2^) and randomly assigned each zone to one of three removal treatments (i.e., aerial, trapping, and toxicant), and a control (i.e., no treatment; Figure 1). We conducted ground shooting efforts within the trapping treatment area, in conjunction with trapping. Overall, we attempted to remove as many wild pigs as possible using each method within its corresponding treatment zone, respectively. We deployed 122 GPS collars on adult wild pigs during Nov 2022–Jan 2023 spread throughout each of the zones for the purposes of another study, thus will not go into details here other than we intentionally did not remove those animals during the aerial and trapping removal efforts (i.e., we did not shoot and released from traps). For the toxicant, we could not control which wild pigs consumed the toxic bait. We did not conduct any removal activities in the control zone.

**Figure 1.**
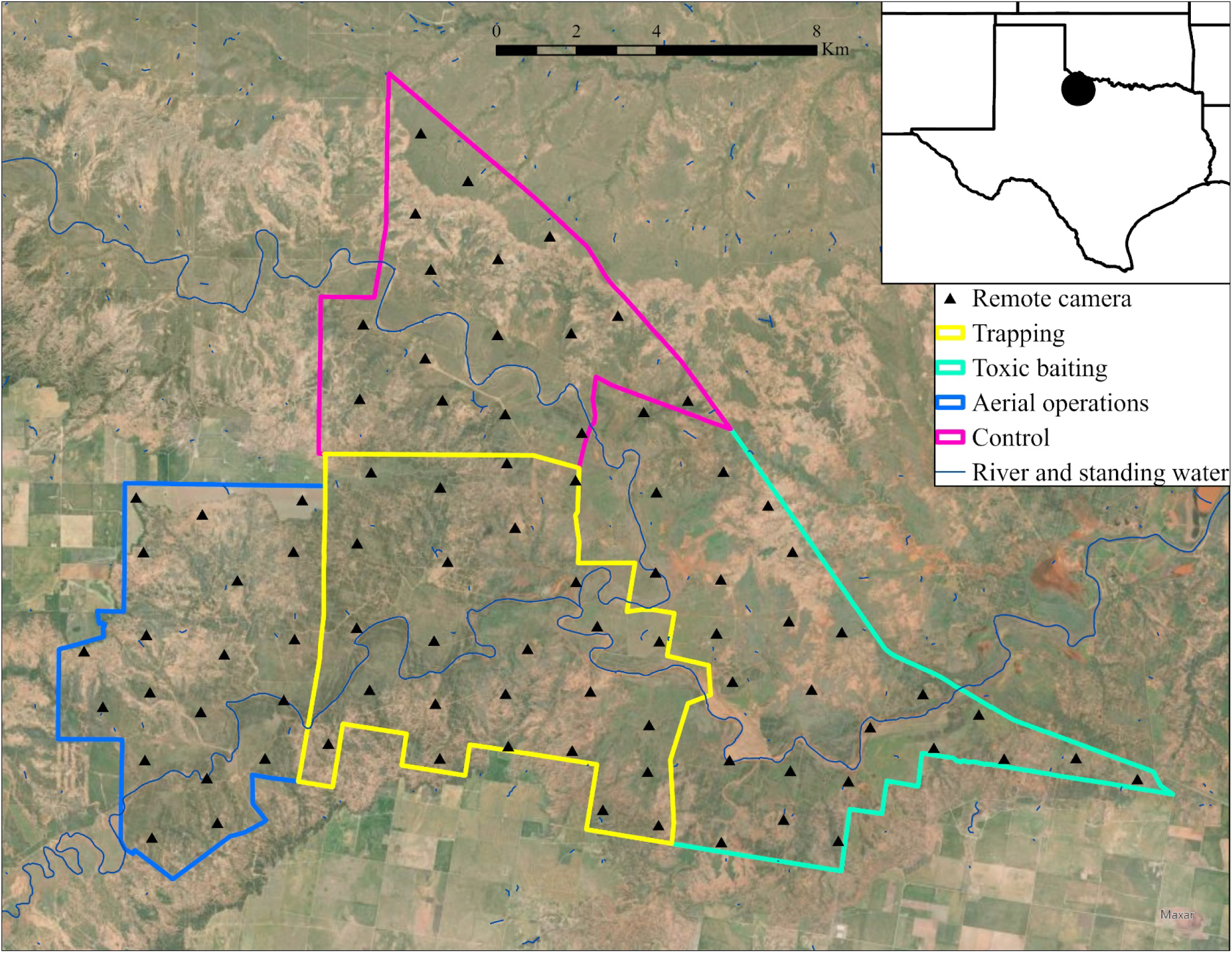
Study area for intensive removals of wild pigs in northcentral Texas, USA during February–April 2023. Remote cameras were used for population estimation.

The timing of control efforts was focused during March–April 2023. Pre-baiting for the trapping and toxicant zones was initiated in mid-Feb 2023. The toxic bait was ultimately deployed during 03–07 March. Trapping and removing wild pigs occurred during 14 March–30 April. Aerial operations for removing wild pigs occurred during 27–29 March. For each method, our goal was to remove the maximum number of wild pigs in the shortest timeframe possible, as would be necessary during an FAD emergency response (see sections below regarding each removal method for more details).

### Population camera grid

We deployed a grid of remote cameras (PC900 or HP2X; RECONYX, Holmen, WI, USA) to evaluate how well our methods reduced the population densities of wild pigs in each of the treatment zones. Prior to deploying the cameras, we recorded the detection area (m) for each camera by recording the maximum distance that the motion-activated trigger for each camera sensed movement as a human walked perpendicular across the field-of-view. The cameras were placed with ∼0° slope. For deployment, we programmed the cameras to record motion-activated images in 5-image bursts with images separated by 3 seconds, and 5-minute quiet periods between bursts.

We generated a camera grid of 85 random locations spread throughout the study area (Figure 1). We attempted to separate sites by ∼2.7 km based on home range sizes from wild pigs nearby (Snow and VerCauteren, 2019) and restricted sites to ≤400 m of a road or clearing to facilitate logistical access. In October 2022, we deployed remote cameras at those sites, ≤50 m from the exact random site based on maximizing the camera field-of-view. Fewer than five random sites were in agricultural fields or open clearings, which we moved ≤300 m to the nearest cover to avoid cameras being damaged or stolen. We set the cameras facing north, ∼1 m off the ground, perpendicular to the ground, and cleared any vegetation that might trigger motion-activated images.

We did not use bait at camera sites. We maintained cameras every ∼2.5 months (i.e., replaced the SD cards and batteries) in attempt to maintain continuous monitoring of sites. Images were initially processed using CameraTrapDetectoR (Burns et al., 2022) for automated species identification of animals (Tabak et al., 2019). Then, we manually verified and labeled all reported wild pig images with a single-observer technique using the Colorado Parks and Wildlife Photo Warehouse Database (Ivan and Newkirk 2016).

### Trapping

We generated a focal area (19.0 km^2^) for intensively trapping wild pigs using the greatest concentration area of GPS collared animals, which we assumed represented the most heavily used portion of the trapping treatment zone (Figure 2). We simultaneously chose a size for our focal area that we estimated we could intensively remove the population of wild pigs considering our limited personnel and resources. Within the focal area, we generated a 500×500 m grid to guide our placement of bait sites, and ultimately the deployment of one trap per grid cell. We chose this size based on wild pig movement patterns relative to bait sites that we measured previously (Davis et al., 2017; McRae et al., 2019; Snow and VerCauteren, 2019). Within each grid cell, we used sign of recent wild pig activity (i.e., rooting, fresh feces, game trails) and landscape features such as ponds, roads, and waterways to place bait sites. We initially deployed ∼20 kg of whole-kernel corn at the sites. We deployed ∼5 kg of corn in a pile near the center of the site, and the remaining corn in the form of 20–50 m corn trail(s) extending in multiple directions. We installed a remote camera (HP2X; RECOYNX) overlooking the central bait pile at each site, set to record motion-activated images in bursts of 3 images (each image separated by 30 seconds), and 5 min quiet periods between bursts. We revisited bait sites every ≤2 days to view camera images and refresh bait as needed. We installed a 3-strand barbed wire fence on steel t-posts around sites, as needed, to exclude cattle from consuming bait. We ensured the lowest strand of barbed wire was ∼0.6 m above ground to exclude young cattle but allow passage by wild pigs.

**Figure 2.**
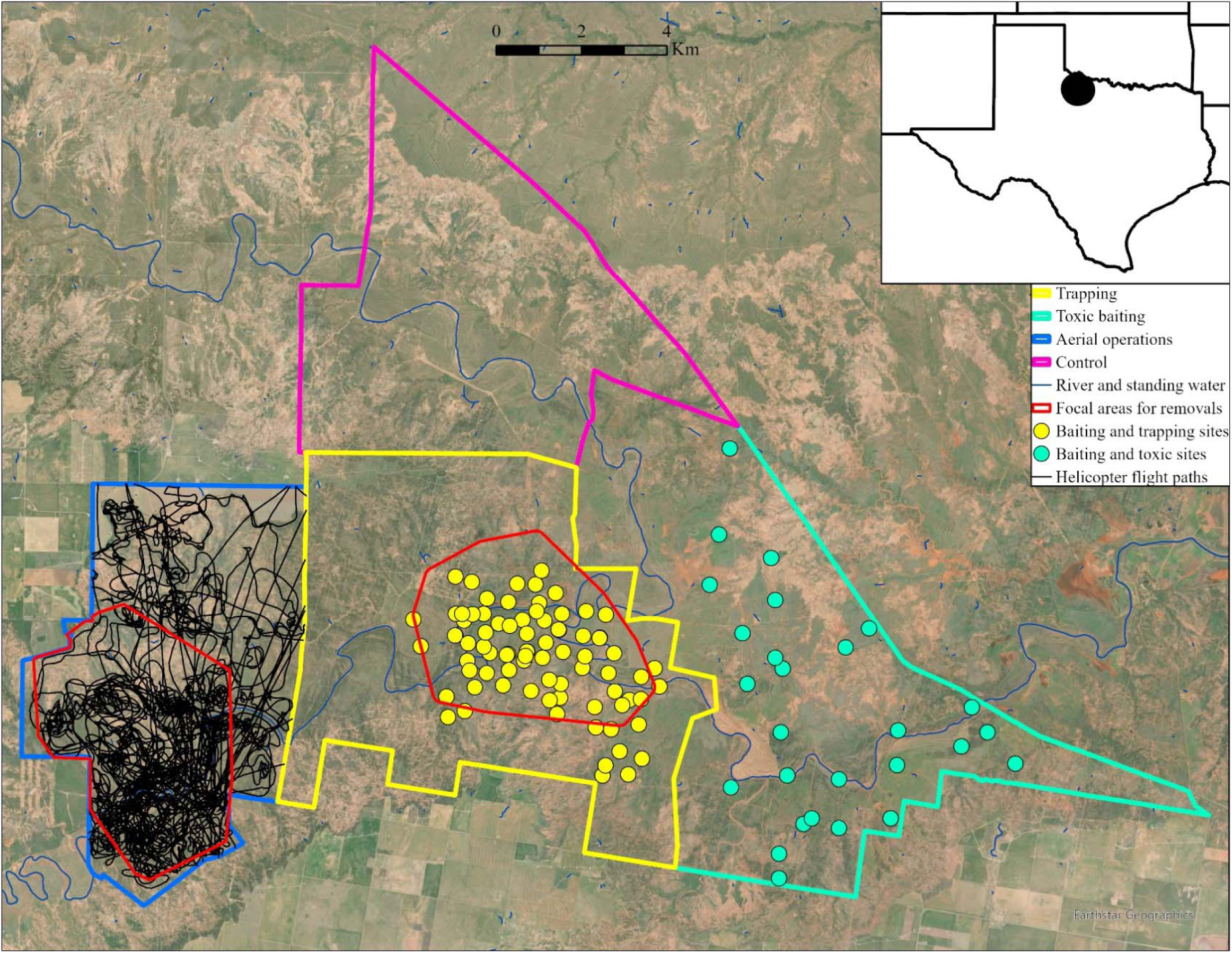
Removal locations and effort for three removal methods for wild pigs in northcentral Texas, USA during February–April 2023. In addition, ground shooting was attempted in the trapping treatment area.

We purposefully used multiple types of traps and deployed each type based on ease of installation at the site, and numbers and behaviors of wild pigs visiting. Overall, we used seven passive net traps (PIG BRIG, Field Engine Wildlife Research & Management, Moodus, CT, USA), nine cellular trap gates with rigid trap panels (M.I.N.E.™ Gate, Jager Pro, Fortson, GA, USA; HogEye camera, latch, and gate system Wildlife Dominion, Crawford, MS, USA), and seven custom-made, collapsible box traps (0.94×0.94×1.65 m). We also had enough extra rigid trap panels for 4-6 additional traps, that we deployed initially without gates and installed the trap gates as gates became available from other trap sites. The passive net traps were a one-way design (i.e., allowing entry with no exit), thus captured wild pigs continuously once set. We primarily used a cellular-activated trigger mechanism on the box traps but also occasionally used an animal-activated trigger (e.g., rooter stick, trip line, snap shackle) in areas without cellular coverage or when cellular-activated triggers were not available. We only installed box traps at sites with single wild pigs visiting. Otherwise, we used net-traps or corral-style traps with rigid panels.

Once wild pigs were observed consistently visiting the bait sites (e.g., ≥2 consecutive nights), we began deploying traps. Our goal was to slowly install the traps piece-by-piece throughout multiple days to slowly condition the wild pigs, but occasionally installed the entire trap at once if deemed more efficient. Our goal was to capture entire groups of wild pigs visiting the trap site (i.e., whole-sounder removal; Lewis et al., 2022). For the passive net traps, we slowly installed and slowly lowered the nets as dictated by wild pig visitation. For the rigid traps and box traps, we slowly installed panels leaving multiple openings for easy access/escape by wild pigs, until routine wild pig visitation allowed closing of all openings except for the trap gate. Once all wild pigs were routinely entering a trap, we set the trap for capture. We monitored the cellular-activated traps in real-time, and remotely closed the trap gates once all wild pigs had entered the traps. For the net traps and animal-activated traps, we returned the following morning to check the traps.

Following any captures, we enshrouded the traps with shade cloth immediately upon arrival to the site (Lavelle et al., 2019). We used a suppressed .22 caliber rifle or pistol to euthanize wild pigs according to AVMA recommendations (Leary et al., 2020). As we removed wild pigs throughout the focal zone, we extended our trapping efforts into adjacent areas that seemed to be additional hotpots of wild pig activity (Figure 2). Ultimately, we pre-baited 78 bait sites and deployed trapping equipment at 66 (84.6%) of those sites. However, we only set traps for capture at 51 (65.3%) of the sites as wild pigs became conditioned. We recorded the total number of wild pigs removed per night per trap from those sites.

### Toxic baiting

For this study, we evaluated an experimental toxic bait containing 5% sodium nitrite (SN; HOGGONE® 2, Animal Control Technologies Australia, Somerton, Victoria, AU), which has been shown to be effective for removing wild pigs (Snow et al., 2017a; Snow et al., 2021b) and low risk of secondary hazards for scavengers of carcasses of wild pigs (Snow et al., 2018; Snow et al., 2019a). Testing of the SN-toxic bait was conducted under an Experimental Use Permit (EUP; 56228-EUP-42) approved by the US Environmental Protection Agency (EPA), with the purposes of evaluating its efficacy and risks to non-target animals. We capitalized on this EUP study to include toxic baiting as a treatment for FAD control, but as such, the study design was slightly different from trapping. Specifically, we spaced toxic sites throughout the toxic treatment zone to ensure independence among the toxic bait sites. Despite not focusing on a smaller focal area, comparisons for time, effort, and costs were representative because our methodologies for deploying the toxic bait simulated operational use.

We implemented a pre-baiting and conditioning strategy that ultimately took ∼10–18 nights to condition and train groups of wild pigs before deploying the SN-toxic bait, as established by previous studies (Lavelle et al., 2018b; Snow et al., 2019b; Snow and VerCauteren, 2019; Snow et al., 2021b). Specifically, we placed and maintained ∼20 kg of whole-kernel corn to pre-bait sites where wild pig activity (e.g., rooting, trails, wallows) was evident for ∼5-7 nights. Then, we transitioned wild pigs to consuming a placebo bait (i.e., identical to the SN-toxic bait but without SN), from within a wild pig-specific bait station designed as a back-to-back trough with lids locked in place by ∼13 kg of magnetic pressure (Snow et al., 2017b; Lavelle et al., 2018a) during the next ∼5–7 days. Once wild pigs were consuming the placebo bait without hesitation, we deployed the SN-toxic bait for one night, followed by 1–2 nights of post-toxic baiting with the placebo bait. If any surviving wild pigs were observed consuming the post-toxic placebo, we deployed the SN-toxic bait for one additional night followed by two nights of post-toxic placebo.

From the onset of pre-baiting, we monitored the sites with remote cameras like described above for trapping. We visited the sites each day to monitor the visitation of wild pigs, refresh bait, and slowly progress toward the ultimate deployment of the SN-toxic bait. We initially deployed 27 bait sites that were ≥500 m apart with the goal of targeting unique groups of wild pigs at each site. Throughout the pre-baiting and acclimation process we narrowed those sites to the best 11 sites for deploying the SN-toxic bait, as restricted by the EUP. We selected the best sites based on the following criteria: (1) consistent visitation by wild pigs, (2) consistent visitation by a family group of wild pigs (i.e., one or more females with multiple juveniles or piglets), and (3) consistent visitation by a family group(s) that were independent from any other family groups observed at other sites. We constructed 3-strand, barbed-wire fences around sites as needed to exclude cattle. At sites with >10 wild pigs visiting, we deployed two bait stations.

Two of the 11 sites did not have wild pigs visit during the night that SN-toxic bait was deployed, therefore only nine sites exposed wild pigs to the SN-toxic bait. We estimated the number of wild pigs removed from consumption of the SN-toxic bait using the remote cameras. Specifically, we counted the unique number of individuals that were observed at the bait sites on the nights that the SN-toxic bait was deployed. Then, we compared these records to the counts of unique individuals observed during the two post-toxic placebo nights. We assumed that any wild pigs that did not return were removed by the SN-toxic bait. In several cases, uniquely identifiable wild pigs visited on post-toxic nights that did not visit during SN-toxic deployment. Therefore, we did not include these individuals in the comparison because they were not exposed to the SN-toxic bait.

### Aerial operations

Similar to the trapping methodologies, we generated a focal area (21.3 km^2^) for intensively removing wild pigs using locational data from wild pigs with GPS collars to identify the most heavily used portion of the aerial treatment zone (Figure 2). We followed strategies outlined by (Davis et al., 2018) and conducted three consecutive days of aerial operations to remove wild pigs. Specifically, we used two helicopters, where the first helicopter was the primary shooting crew, and the second helicopter was a spotting and chasing crew. Each helicopter crew consisted of a pilot and gunner. The gunners used a combination of semiautomatic rifles and shotguns to remove wild pigs. Helicopters worked together and attempted to remove all wild pigs from within a group. As densities of wild pigs were reduced in the focal zone, the helicopters extended outside of the focal zone (but still within the aerial treatment zone) to conduct removals of adjacent wild pigs. We did not count the adjacent wild pigs removed or the time spent removing of recovering wild pigs because we did not attempt to locate and recover those carcasses.

We recorded the total numbers of wild pigs removed based on counts from the helicopter crews. In some cases, it was difficult for the helicopter crews to determine the exact number of wild pigs they removed from within a group, especially with large groups of small wild pigs. Therefore, they estimated the number to the best of their ability or occasionally reported “multiple piglets”. We conservatively considered reports of multiple piglets to be five piglets, although recognize that groups with ≥5 piglets were commonly observed in our study area. We chose five because we assumed 1–4 piglets would likely have been counted, but groups ≥5 were more difficult to quickly count from the air.

### Ground shooting

Ground shooting was not a primary removal method and attempted only in a few situations in the trapping zone. First, we attempted to shoot wild pigs that were observed outside of traps, while checking traps, only if it was deemed we would not educate the animals from visiting the traps again if unsuccessful (e.g., only 1 wild pig present). Second, occasionally in the trapping zone we observed bait sites with only a single wild pig visiting. Instead of moving a trap to that site, we would attempt to ground shoot the wild pig at night using rifles equipped with thermal or night vision optics. Finally, occasionally we baited the 2-track roads with whole-kernel corn and attempted to shoot wild pigs at night that were on the roads. All ground shooting was only attempted as availability of personnel hours allowed.

### Carcass recovery

We attempted to locate and mock-recover every carcass from all four removal methods. We did not remove or dispose of the carcasses as would be required during an emergency response. Measurements were collected from recovered carcasses and remote cameras were deployed on a subset of the carcasses for the purposes of another study. We assumed the time required to take measurements and deploy cameras was comparable to the time required to load and dispose of the carcasses during a disease response. We recorded sex, age from tooth eruption (Halseth et al., 2018), weight (kg), body dimensions, and GPS location for any found carcasses. For the toxic treatment, we also examined for evidence of SN-toxicity by visual inspection of the darkness of a drop of blood on a white laminated card compared to a standard curve representing the percentage of methemoglobin in the blood (Patton et al., 2016). All time spent locating and recovering carcasses were considered as carcass recovery hours.

For trapping, recovery of carcasses was straightforward. We removed the carcasses from the traps and brought them to disposal locations that would not interfere with future trapping efforts (i.e., >200 m from any trap site). To transport carcasses, we loaded them into the back of off-highway vehicles (OHVs) or dragged them using a rope. Carcasses from ground shooting were similarly recovered immediately at the location they were removed.

For the toxic baiting, we conducted systematic searches for carcasses of wild pigs for two consecutive days following each deployment of SN-toxic bait, following Snow et al. (2021b). Specifically, we generated a 400×400 m grid with transects spaced every 50 m centered over each bait site. We uploaded georeferenced maps of the transects onto handheld devices, and used our real-time locations to follow the transects and conduct the searches. We walked north-south transects on the first morning, and east-west transects on the second morning following deployment of SN-toxic bait. We did not walk transects if no animals accessed the bait stations on the nights that SN-toxic bait was deployed. Typically, 2–4 people walked the transects for each bait station. Once a dead wild pig was located, we searched the nearby area for other carcasses even if it meant leaving the grid, because wild pigs often feed and die in groups (Snow et al., 2021b). If carcasses were not located within grids, but wild pigs were suspected to have consumed enough toxic bait to die, we conducted extra searching outside of the transects by driving and scanning/smelling for decomposing carcasses, by searching for turkey vultures (*Cathartes aura*), or listening for groups of coyotes (*Canis latrans*) that might be scavenging on carcasses. We also followed any fresh tracks from wild pigs on game trails off the grids as possible.

For the aerial operation, we devised a system of communication between the helicopter crews and carcass recovery crews. The helicopter crews radioed coordinates of each carcass to a centralized dispatch person on the ground. The dispatch person compiled locations of 1–10 wild pigs that were in proximity to each other and dispatched a carcass recovery crew to those groups of carcasses, prioritizing the crews that were currently closest. Carcass recovery crews consisted of two people on OHVs, of which we had 5 crews. The crews navigated to the carcasses using handheld GPS devices. Ultimately the helicopter crews removed wild pigs faster than the carcass recovery crews could locate the carcasses, therefore carcass recovery proceeded for hours after the helicopters were done shooting, and in some cases the day after.

### Time and effort data collection

We generated real-time maps of bait and carcass sites, and used these maps for navigation using ArcGIS Collector (ESRI, Redlands, CA USA) on mobile devices (i.e., tablets or smart phones). We also collected information on activities for the trapping and toxic baiting efforts from each person that was conducting these activities using ArcGIS Survey123 (ESRI, Redlands, CA USA). Trapping activities included: camera check, pre-bait site, build trap, set trap, capture and remove pigs, trap removal, and other (that we specified in a comment). Toxic baiting activities included: camera check, pre-bait site, build bait site cattle-fence, deploy bait stations, toxic baiting application, toxic baiting clean up, bait site removal, and other. For both, we also recorded the date bait sites were established, when and what was constructed (e.g., trap type or bait station), how much and type of bait deployed daily, and the number and morphometric data associated with every wild pig removed. We recorded the latter for the aerial and ground shooting operations as well.

We also recorded the personnel hours, per individual, needed to complete certain tasks (hrs rounded to the nearest 0.5 hr) using ArcGIS Survey123. Trapping categories included: building/setting/checking traps, cellular trap monitoring, tear down trap, and euthanizing and carcass management. Toxic baiting categories included: establish/check pre-bait sites, building/baiting/checking bait sites, toxic baiting clean up, bait site removal/relocation, and carcass searching. Aerial operations categories included: ground crew duties (e.g., fueling and communication), pilot and gunner duties, and carcass searching. Ground shooting hours included time spent shooting. Finally, we included a category for time spent conducting logistical duties not related to any specific removal treatment, which included purchasing supplies (e.g., bait), moving and organizing equipment, repairing flat tires, etc.

### Establishing costs

We compiled two lists of costs for each method of removal. The first was a list of costs that were required to operate the removal efforts. These costs included the personnel hours of all individuals involved, costs for bait and other disposables, and any supplies that would be unlikely to be used again (e.g., barbed wire or steel t-posts). We used this list to calculate a cost/wild pig ($USD 2024) for removal and carcass recovery, assuming all the initial investments in equipment were already made. The SN-toxic bait and associated supplies were not available in the USA, therefore we used current pricing from AU and converted to $USD (https://cdn.environment.sa.gov.au/landscape/docs/ep/PestControlPricingSchedule-Jan2023.pdf). We considered the hourly rates for all personnel hours at $19.86/hr, and $29.81/hr for overtime hours (i.e., more than 40 hrs/week per person) which was equivalent to a GS 6–Step 5 Wildlife Specialist for USDA/APHIS/Wildlife Services in 2024. Hourly rates for the helicopter pilot and gunner were included in the overall helicopter cost/hr, thus were not considered separately for that method. The helicopter cost/hr also included costs for operation and for required maintenance. For all methods, we did not include transportation costs to-and-from the study site (i.e., trucks, fuel, and personnel hours). We also did not include costs for repairing vehicles (i.e., trucks and OHVs) that broke during our work, other than the time spent conducting logistical duties (e.g., recovering broken vehicles from the field, fixing flat tires), but we recognize that repair costs were substantial in our study and led to some missed opportunity costs (e.g., necessity of repairing vehicles before could resume baiting wild pigs), thus should be considered for FAD responses. Lastly, we did not include fuel costs for OHVs.

The second was a list of costs for the initial investment in equipment needed to conduct the removals. Some equipment could be reused from previous projects (e.g., traps, bait stations, helicopters, firearms, etc.), but we generated costs as if buying new for a consistent comparison. The one exception to buying new was pre-owned helicopters for practicality. However, the prices varied widely (i.e., $128,000–599,000) based on the hours of operation on various components, therefore we averaged the costs of multiple pre-owned helicopters that would be sufficient for operational use. We did not include costs of buying vehicles (i.e., trucks, trailers, and OHVs) because these would likely already be purchased and the need would vary based on location.

### Data analysis

#### Time and cost comparison

For the time-spent comparison, we summed the hours spent on all removal and logistical activities. Then, for each removal activity, we divided the cumulative hours by the total number of wild pigs removed to calculate a rate of wild pigs removed/person hour. Finally, within each removal activity we calculated the division of hours spent on specific tasks.

For the cost comparison, we summed the total operational costs (i.e., not including initial investment) for each removal activity. We divided the cumulative operational costs by the total number of wild pigs removed to calculate a cost/wild pig. Finally, we used descriptive statistics to compare the total initial investments for each treatment type.

#### Change in population density

We estimated changes in wild pig density for each treatment area pre-versus post-removal using time-to-event models (Moeller et al., 2018) using the spaceNtime package in Program R (Moeller and Lukacs, 2022). The time-to-event model uses repeated measures of the time until an event (i.e., wild pig captured on camera), an estimate of animal movement speed, and the detection distance of each camera to estimate population density (Jennelle et al., 2002; Parsons et al., 2017). To determine the change in density relative to removal activities, we restricted the use of camera data to only those cameras that were located ≤ 1,000 m of a trap site, toxic bait site, or helicopter flight path. For the control area, all cameras >1,000 m from any of the removal areas were used. Ultimately, 70 camera sites (aerial = 15, toxicant = 23, trapping = 16, control = 16) with 63,141 trigger events met the criteria for population density estimation.

We used the detection distances from each remote camera and speed of travel of wild pigs to parameterize the time-to-event model. For speed of travel, we estimated the average speed (m/sec) of wild pigs from the GPS collars across the study site, assuming straight-line distances among consecutive 15–20 minute fixes. We determined the movement rates were 0.02 m/sec pre-removal and 0.03 m/sec post-removals.

Estimates of density were generated for 45 days pre- and post-removal and for the trapping zone density was also estimated during the removal period. Control area density was estimated for the period 45 days before the first treatment (aerial operations) and 45 days after the end of the last treatment (trapping). Abundance was estimated by scaling the density up to the area treated. Reliability of model estimates were evaluated by estimating the absolute change in abundance using pre- and post-removal abundance estimates and comparing to the known number of animals removed in each area. We made relative comparisons on population changes among the removal techniques by calculating how much time and costs were required to reduce the estimated pre-removal densities by 10% for each removal type. This relative comparison allows accounting for local differences in initial densities of wild pigs to enhance our understanding of time- and cost-efficiencies. We acknowledge that time and costs to reduce densities increase as densities are reduced (e.g., Fischer et al., 2020), therefore our inferences are relative to a rapid reduction in the initial population and not elimination.

#### Recovery of carcasses

For the toxic baiting and aerial operations methods, which both included carcass searching time, we calculated the rates recovery we achieved (carcasses /km^2^/person hour). Specifically, for toxic baiting we calculated the area searched as 0.16 km^2^ per grid × 9 grids × 2 days = 2.88 km^2^. For the aerial operations we used the full area of the focal zone as the area searched (21.3 km^2^), although we had coordinates of carcasses to direct our search.

Finally, we compared the distributions of ages of wild pigs removed and recovered among each method using descriptive statistics. For each age class from Halseth et al. (2018), we calculated a median age each wild pig. For example, if we aged a wild pig in the 20–30 week age class, we estimated it was 25 weeks (i.e., 5.75 months). We plotted the distributions of ages for comparison amongst removal methods. We did not include ground shooting because too few animals were removed using this method.

## Results

Overall, we removed 612 wild pigs, including 296 with trapping, an estimated 58 with toxic baiting, an estimated 256 with aerial operations, and 2 with ground shooting (Table 1). The overall duration of the trapping effort (i.e., including pre-baiting and building) was 73 days and included ∼728 total trap nights (i.e., sum of traps per night that were set and ready to catch wild pigs). The duration of toxic baiting was 23 days and limited to 15 total toxic bait nights (i.e., sum of bait sites per night that had toxic bait deployed). The duration of the aerial operation was three days, which totaled six helicopter days (i.e., sum of helicopters per day). The aerial operation removed an additional ∼441 wild pigs from areas adjacent to our focal zone which we did not consider in our counts. Finally, ground shooting occurred throughout a 47-day period and included 18 shooting nights (i.e., sum of people per night that were actively hunting wild pigs).

**Table 1.** Summary of removal efforts for wild pigs in northcentral Texas, USA during February–April 2023.

| | Area<br>(km<sup>2</sup>) | Duration<br>(days) | Total<br>personnel<br>hours | Total<br>operational<br>costs<br>(\$USD) | Initial<br>investment<br>costs<br>(\$USD) | Wild pigs<br>removed | Carcasses<br>recovered | Change in<br>density pre-<br>vs. post-removals<br>(%) |
| --- | --- | --- | --- | --- | --- | --- | --- | --- |
| Trapping | 19.0 | 73 | 2,055 | \$72,797 | \$125,198 | 296 | 296 | -89.9 |
| Toxic baiting | 64.9 | 23 | 656 | \$30,711 | \$33,156 | 58 | 43 | -23.3 |
| Aerial operation | 21.3 | 3 | 298 | \$37,623 | \$772,780 | 256 | 126 | -51.8 |
| Ground shooting | 19.0 | 18 | 53 | \$1,417 | \$21,847 | 2 | 2 | NA |
| Total | 124.2 | 117 | 3062 | \$142,548 | \$952,981 | 612 | 467 | -55.0 (mean) |

### Time and cost comparison

Overall, we spent 3,964 person hours to remove 612 wild pigs and recover their carcasses, averaging 0.15 wild pigs/person hour among all four removal methods, excluding logistical hours. We found that the aerial operation was 9.1 times more efficient per hour at removing wild pigs and recovering carcasses than the next most efficient method (Figure 3). The next most time efficient methods were trapping, then toxic baiting, and then ground shooting. Cumulatively, we spent nearly a quarter of our time on logistical activities shared across all removal methods (Figure 4). Nearly 75% of the hours for trapping were spent pre-baiting or building traps, including substantial time spent building exclusion fencing for cattle. Whereas the labor for setting up toxic baiting was less (i.e., 57% for pre-baiting and building), and would have been similarly less had cattle not been present. Carcass recovery also took a substantial amount of time with the toxic bait (i.e., 27%). The most time intensive time-component of the aerial operation was carcass recovery (i.e., 59%).

**Figure 3.**
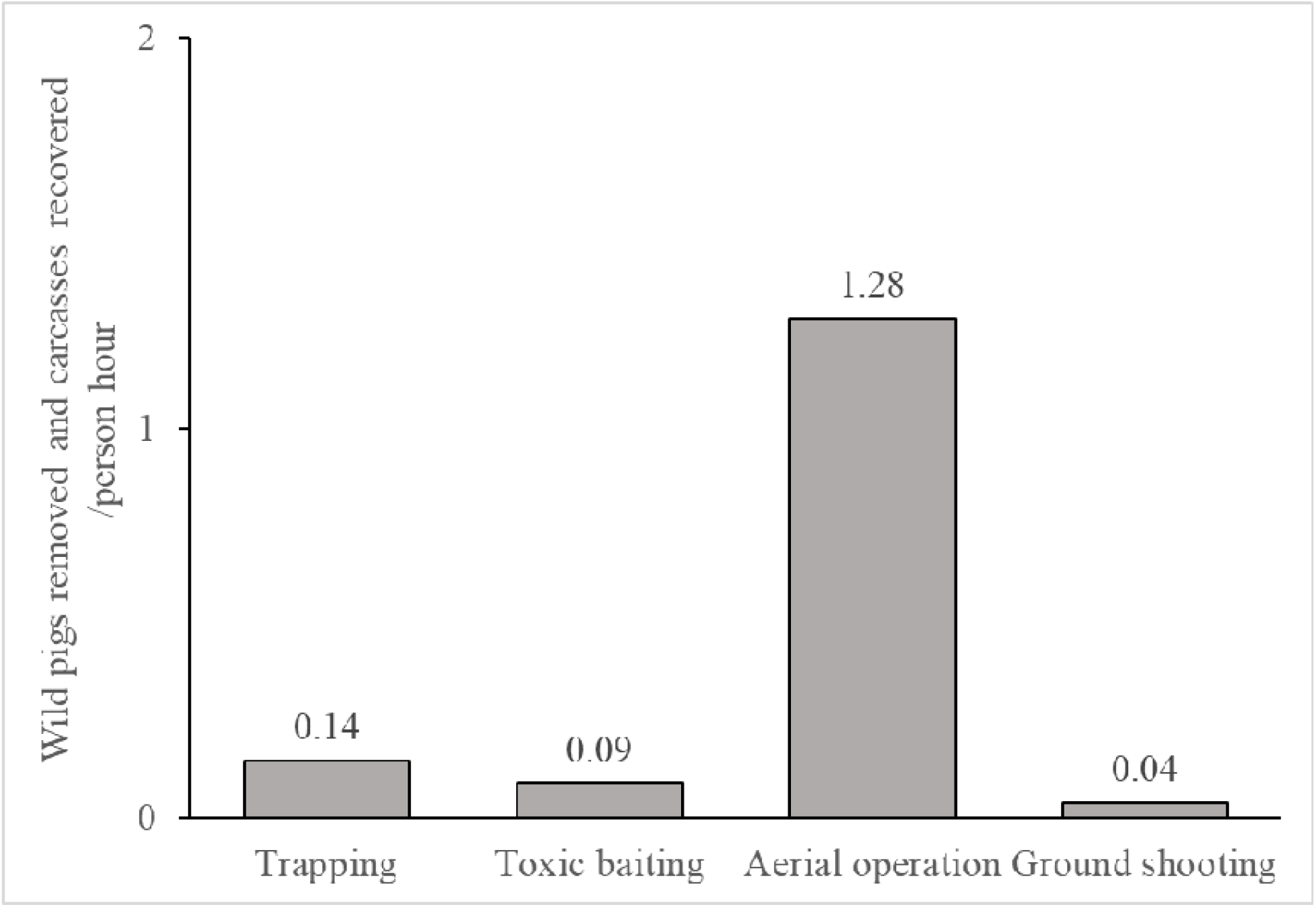
Comparison of wild pigs removed and carcasses recovered/person hour for each removal method for removing and recovering carcasses of wild pigs in northcentral Texas, USA during February–April 2023. Results exclude logistical hours which were shared among the methods.

**Figure 4.**
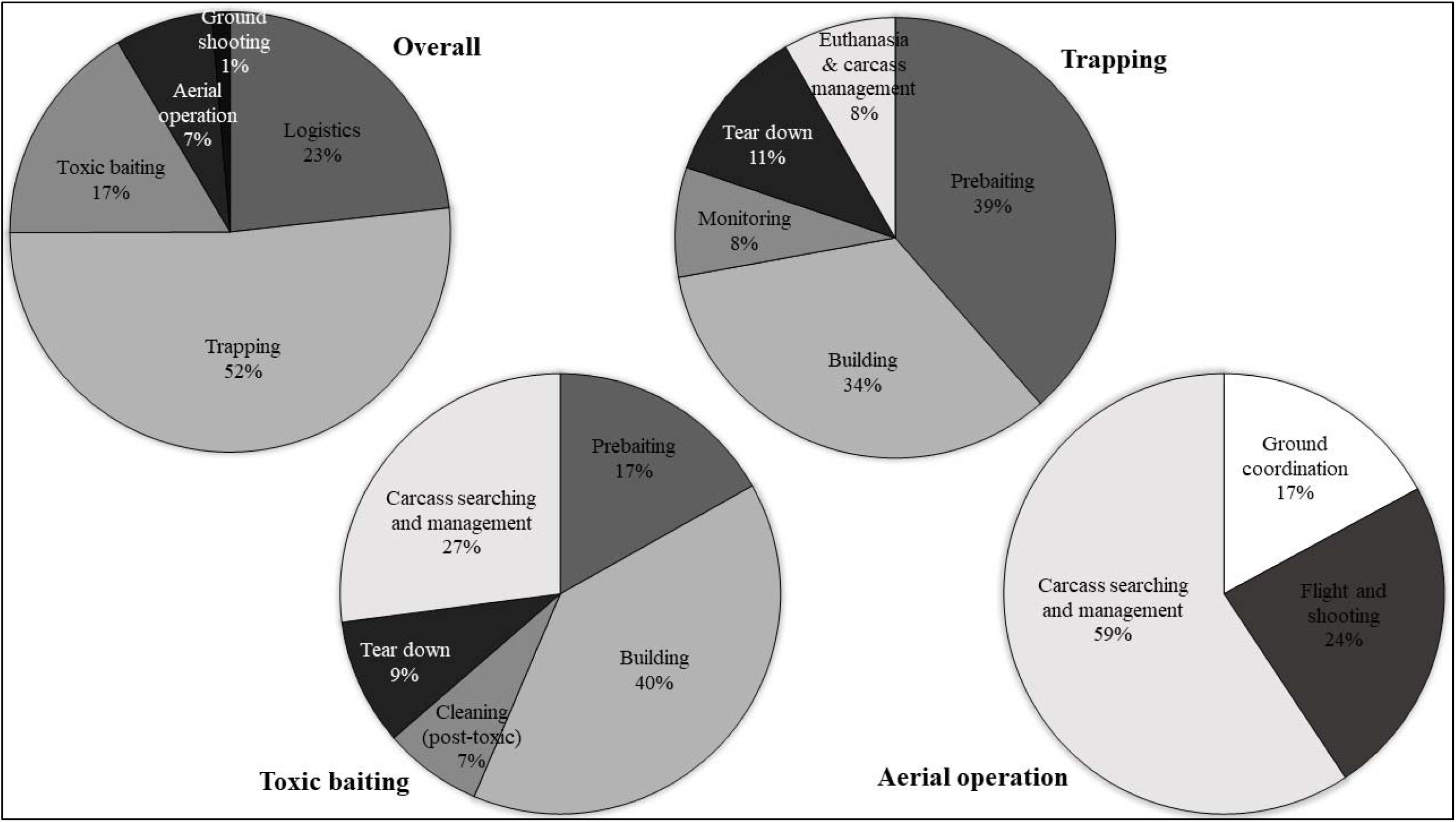
Distribution of time spent (hours worked) during intensive removal activities for wild pigs in northcentral Texas, USA during February–April 2023. Cumulatively, 3,964 hours were worked (logistics = 922, trapping = 2049, toxic baiting = 656, aerial operation = 284, ground shooting = 54) and 612 wild pigs were removed (trapping = 296, toxic baiting = 58 [estimated], aerial operation = 256, ground shooting = 2). Distribution for ground shooting hours excluded below because they were 100% hunting.

Cumulatively, we spent $142,623 in operational costs to remove and recover carcasses of 612 wild pigs, averaging $233.04/wild pig (Table 2). The most cost-effective method for removing and recovering carcasses was the aerial operation ($147.26/wild pig). Trapping was 1.67 times more expensive, toxic baiting was 3.60 times more expensive, and ground shooting was 4.81 times more expensive per wild pig.

**Table 2.**
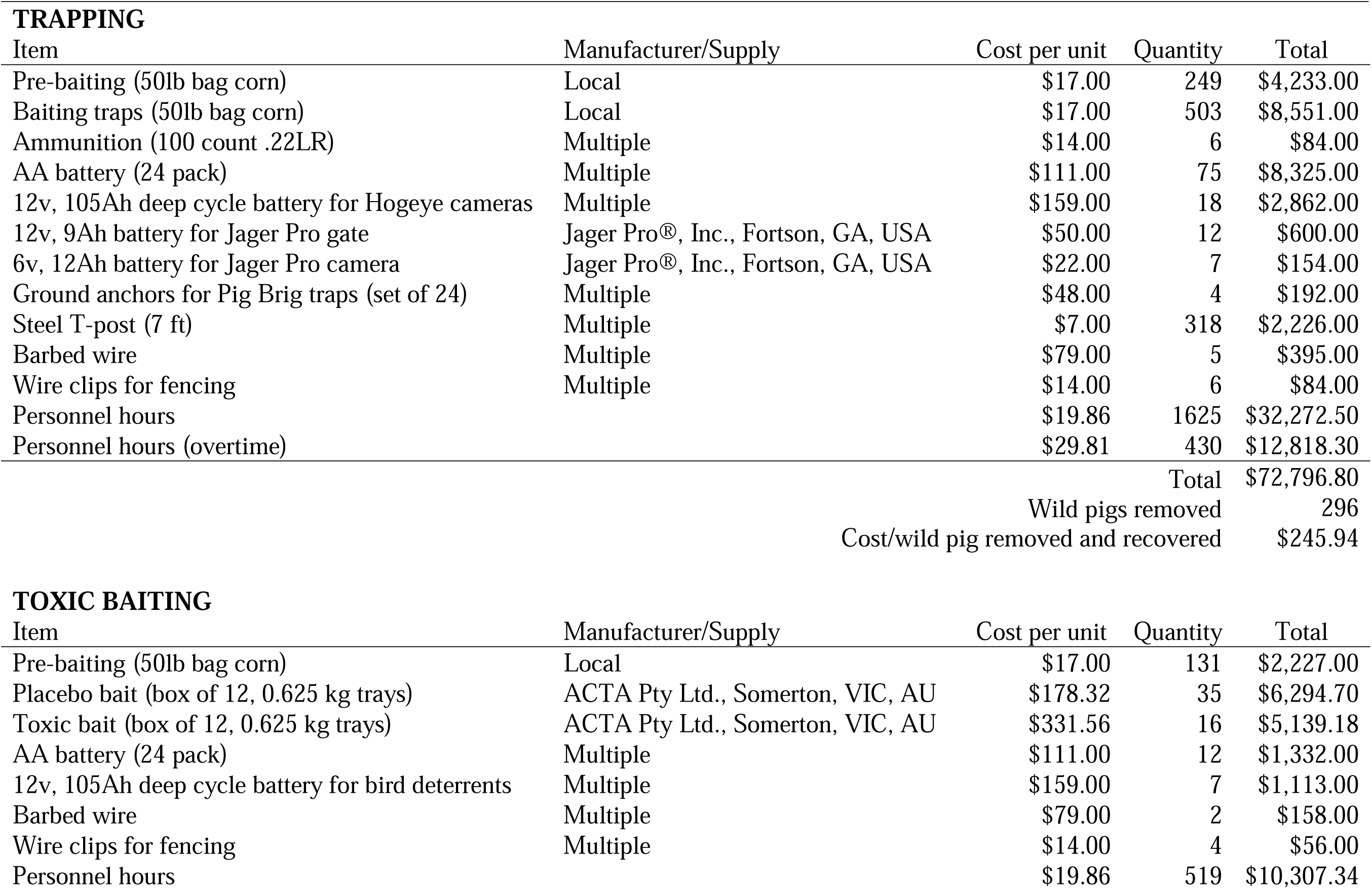
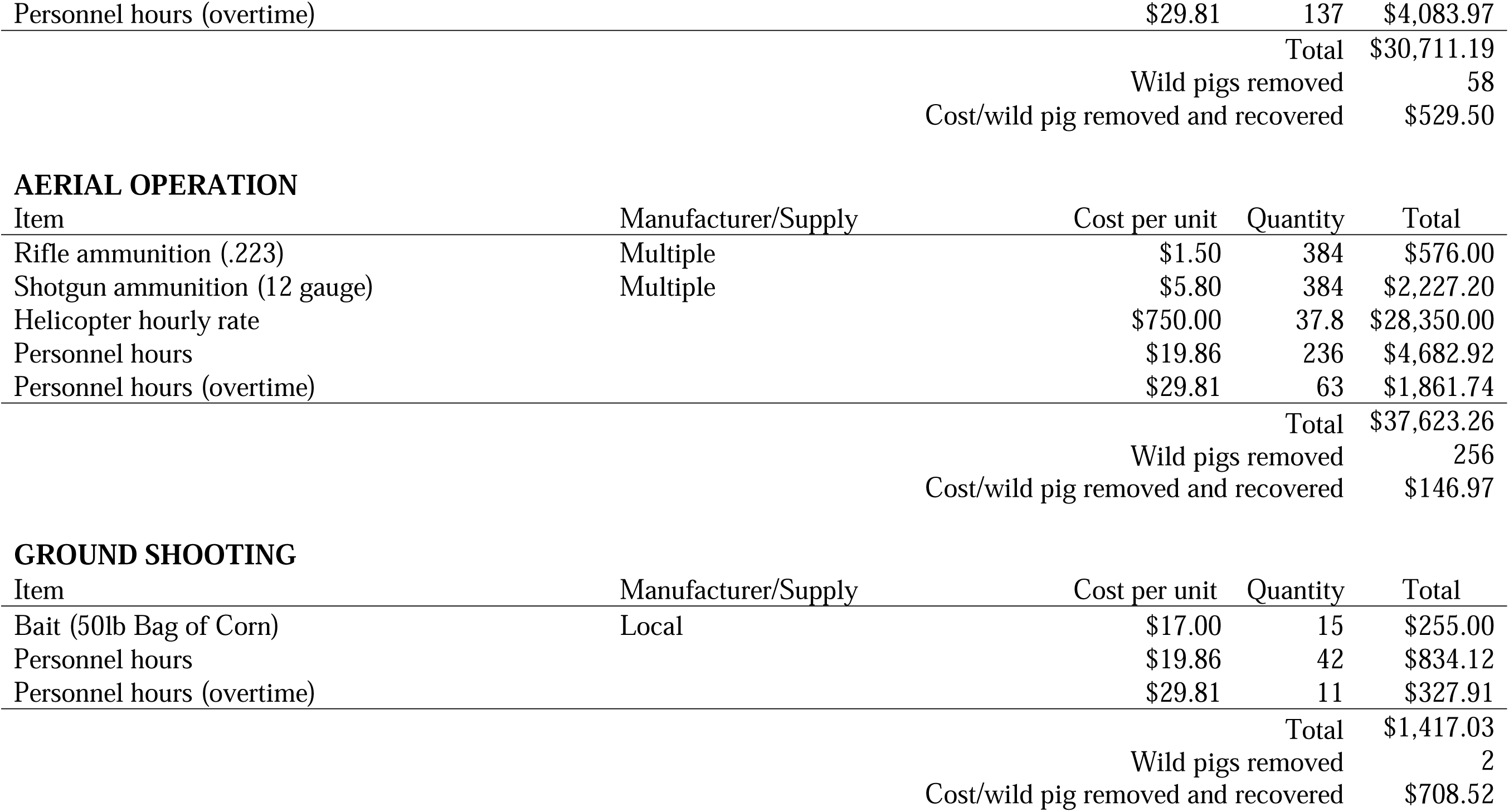
Estimates of operational costs including personnel hours, bait and other disposables, and any supplies that would be unlikely to be used again for removal methods for wild pigs in northcentral Texas, USA during February–April 2023.

We estimated the initial investment into equipment was $952,981 among all four methods used in our study (Table 3). Aerial operations was the most expensive and comprised 81% of that total initial investment. Trapping required the second most initial investment but was 84% less expensive than aerial operations. Toxic baiting was third and required 96% less initial investment than aerial operations. Ground shooting required the least amount of initial investment and required 97% less initial investment than aerial operations.

**Table 3.**
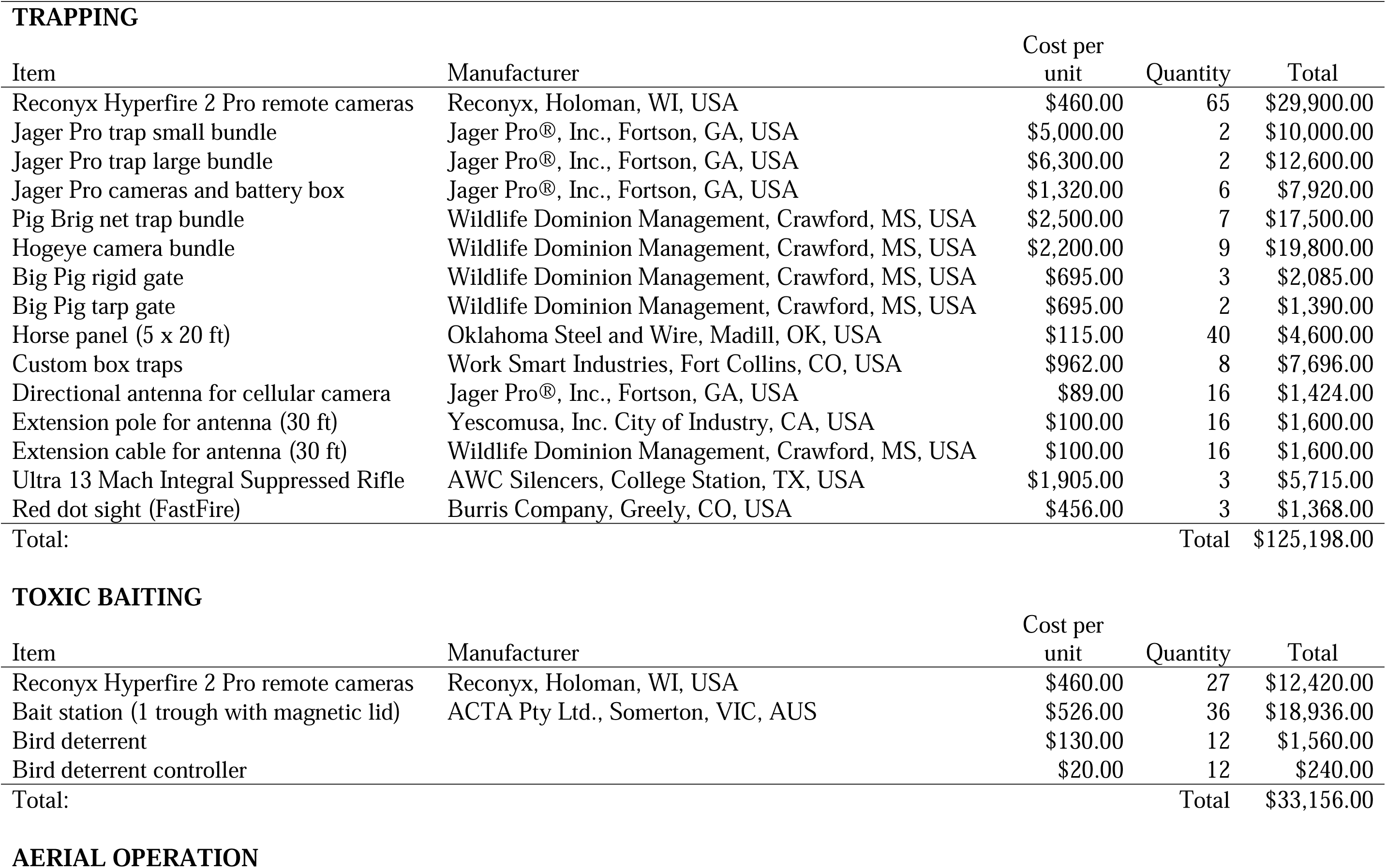
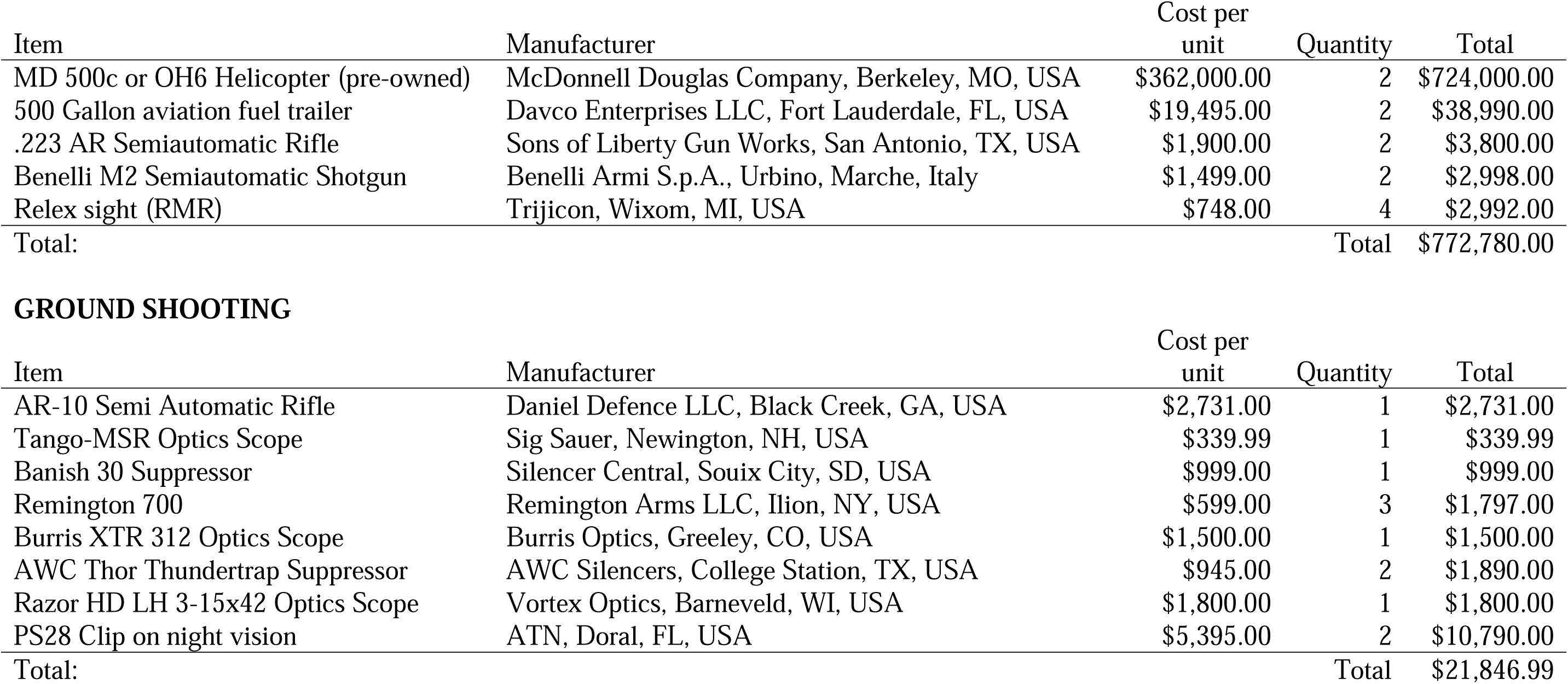
Estimates of initial equipment investment for removal methods for wild pigs in northcentral Texas, USA during February– April 2023.

| **TRAPPING** |  |  |  |  |
| --- | --- | --- | --- | --- |
| Item | Manufacturer | Cost per unit | Quantity | Total |
| Reconyx Hyperfire 2 Pro remote cameras | Reconyx, Holoman, WI, USA | \$460.00 | 65 | \$29,900.00 |
| Jager Pro trap small bundle | Jager Pro®, Inc., Fortson, GA, USA | \$5,000.00 | 2 | \$10,000.00 |
| Jager Pro trap large bundle | Jager Pro®, Inc., Fortson, GA, USA | \$6,300.00 | 2 | \$12,600.00 |
| Jager Pro cameras and battery box | Jager Pro®, Inc., Fortson, GA, USA | \$1,320.00 | 6 | \$7,920.00 |
| Pig Brig net trap bundle | Wildlife Dominion Management, Crawford, MS, USA | \$2,500.00 | 7 | \$17,500.00 |
| Hogeye camera bundle | Wildlife Dominion Management, Crawford, MS, USA | \$2,200.00 | 9 | \$19,800.00 |
| Big Pig rigid gate | Wildlife Dominion Management, Crawford, MS, USA | \$695.00 | 3 | \$2,085.00 |
| Big Pig tarp gate | Wildlife Dominion Management, Crawford, MS, USA | \$695.00 | 2 | \$1,390.00 |
| Horse panel (5 x 20 ft) | Oklahoma Steel and Wire, Madill, OK, USA | \$115.00 | 40 | \$4,600.00 |
| Custom box traps | Work Smart Industries, Fort Collins, CO, USA | \$962.00 | 8 | \$7,696.00 |
| Directional antenna for cellular camera | Jager Pro®, Inc., Fortson, GA, USA | \$89.00 | 16 | \$1,424.00 |
| Extension pole for antenna (30 ft) | Yescomusa, Inc. City of Industry, CA, USA | \$100.00 | 16 | \$1,600.00 |
| Extension cable for antenna (30 ft) | Wildlife Dominion Management, Crawford, MS, USA | \$100.00 | 16 | \$1,600.00 |
| Ultra 13 Mach Integral Suppressed Rifle | AWC Silencers, College Station, TX, USA | \$1,905.00 | 3 | \$5,715.00 |
| Red dot sight (FastFire) | Burris Company, Greely, CO, USA | \$456.00 | 3 | \$1,368.00 |
| Total: | | | Total | \$125,198.00 |
| **TOXIC BAITING** |  |  |  |  |
| Item | Manufacturer | Cost per unit | Quantity | Total |
| Reconyx Hyperfire 2 Pro remote cameras | Reconyx, Holoman, WI, USA | \$460.00 | 27 | \$12,420.00 |
| Bait station (1 trough with magnetic lid) | ACTA Pty Ltd., Somerton, VIC, AUS | \$526.00 | 36 | \$18,936.00 |
| Bird deterrent | | \$130.00 | 12 | \$1,560.00 |
| Bird deterrent controller | | \$20.00 | 12 | \$240.00 |
| Total: | | | Total | \$33,156.00 |
| **AERIAL OPERATION** |  |  |  |  |

| Item | Manufacturer | Cost per unit | Quantity | Total |
| --- | --- | --- | --- | --- |
| MD 500c or OH6 Helicopter (pre-owned) | McDonnell Douglas Company, Berkeley, MO, USA | \$362,000.00 | 2 | \$724,000.00 |
| 500 Gallon aviation fuel trailer | Davco Enterprises LLC, Fort Lauderdale, FL, USA | \$19,495.00 | 2 | \$38,990.00 |
| .223 AR Semiautomatic Rifle | Sons of Liberty Gun Works, San Antonio, TX, USA | \$1,900.00 | 2 | \$3,800.00 |
| Benelli M2 Semiautomatic Shotgun | Benelli Armi S.p.A., Urbino, Marche, Italy | \$1,499.00 | 2 | \$2,998.00 |
| Relex sight (RMR) | Trijicon, Wixom, MI, USA | \$748.00 | 4 | \$2,992.00 |
| Total: | | | Total | \$772,780.00 |

GROUND SHOOTING
| Item | Manufacturer | Cost per unit | Quantity | Total |
| --- | --- | --- | --- | --- |
| AR-10 Semi Automatic Rifle | Daniel Defence LLC, Black Creek, GA, USA | \$2,731.00 | 1 | \$2,731.00 |
| Tango-MSR Optics Scope | Sig Sauer, Newington, NH, USA | \$339.99 | 1 | \$339.99 |
| Banish 30 Suppressor | Silencer Central, Souix City, SD, USA | \$999.00 | 1 | \$999.00 |
| Remington 700 | Remington Arms LLC, Ilion, NY, USA | \$599.00 | 3 | \$1,797.00 |
| Burris XTR 312 Optics Scope | Burris Optics, Greeley, CO, USA | \$1,500.00 | 1 | \$1,500.00 |
| AWC Thor Thundertrap Suppressor | AWC Silencers, College Station, TX, USA | \$945.00 | 2 | \$1,890.00 |
| Razor HD LH 3-15x42 Optics Scope | Vortex Optics, Barneveld, WI, USA | \$1,800.00 | 1 | \$1,800.00 |
| PS28 Clip on night vision | ATN, Doral, FL, USA | \$5,395.00 | 2 | \$10,790.00 |
| Total: | | | Total | \$21,846.99 |

### Change in population density

For the time-to-event model for estimating population densities, we found the population densities declined 89.9% in the trapping zone following the removal period, declined 23.3% in the toxic baiting zone, and declined 51.8% in the aerial operations zone (Table 1, Figure 5, Supplemental Table 1). These estimates of declines were within ≤1% of the actual number of wild pigs we removed in the aerial and toxicant zones, and 6.6% in the trapping zone, indicating these were reliable estimates (Supplemental Figure 1). We did not observe a change in population densities pre- versus post-removals in the control zone. Relative comparisons to reduce pre-removal densities by 10% found that trapping and aerial operations had similar costs, $8,101 and $7,266 respectively (Table 4). Toxic baiting was the most expensive at $13,192 per 10% reduction in population density. The number of personnel hours required for a 10% reduction in population density were lowest for aerial operations, and similar between trapping and toxic baiting.

**Figure 5.**
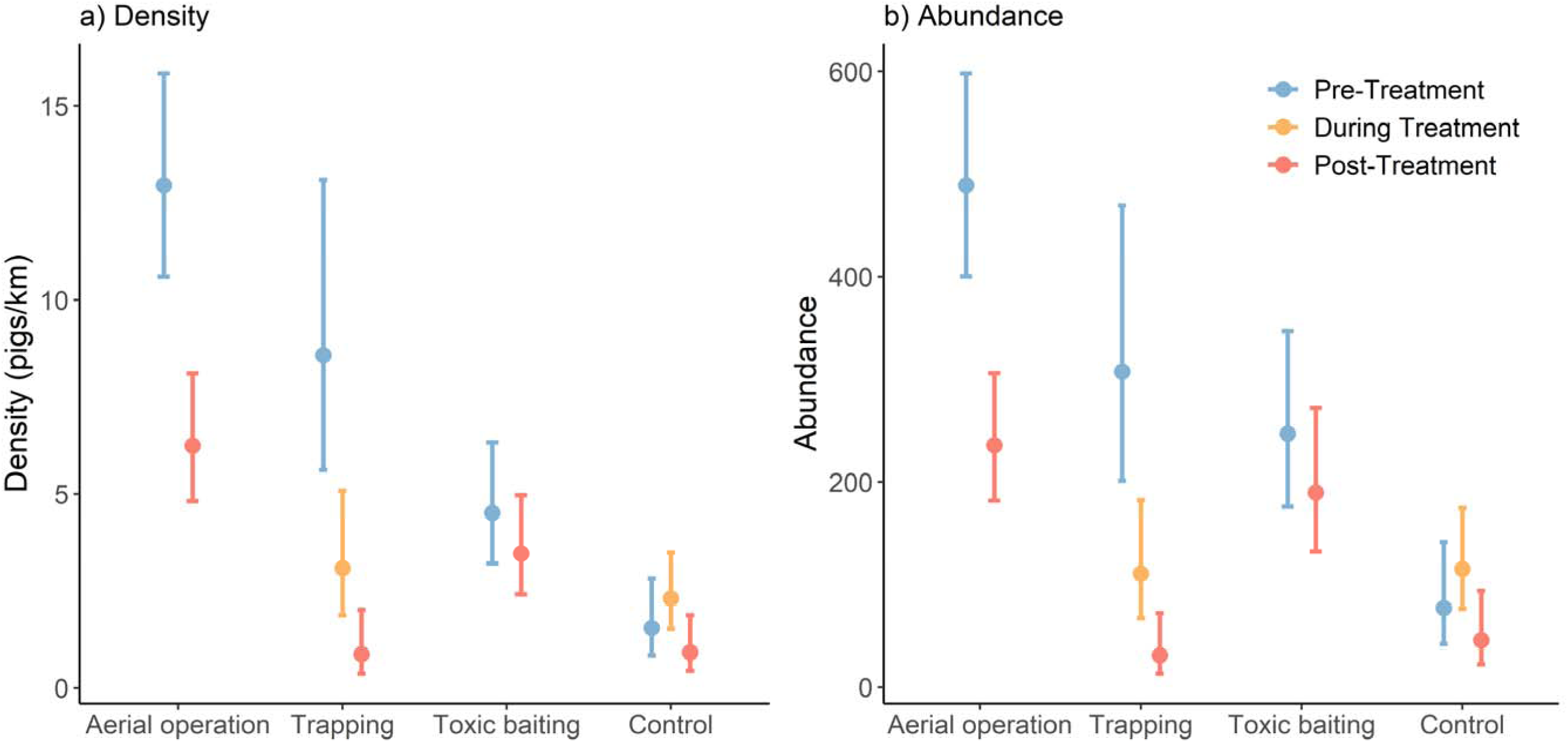
Estimated density and abundance pre-, during, and post-removals of wild pigs in northcentral Texas, USA during February– April 2023. Points indicate estimated mean density and abundance, respectively, with 95% CIs.

**Table 4.** Estimates of operational costs ($USD 2023), personnel hours, and days required to reduce wild pig density by 10% by method in northcentral Texas, USA during February–April

|  | Operational<br>costs | Personnel<br>hours | Days<br>required |
| --- | --- | --- | --- |
| Trapping | \$8,101 | 229 | 8.1 |
| Toxic baiting | \$13,192 | 282 | 9.9 |
| Aerial operation | \$7,266 | 58 | 0.6 |

### Recovery of carcasses

For recovery of carcasses, we were able recover 100% of the carcasses from trapping and ground shooting, respectively. We spent 170 hours euthanizing and managing carcasses during trapping. For toxic baiting, we spent 117 person hours and recovered 43 of the estimated 58 carcasses (72.4%); a recovery rate of 0.08 carcasses/km^2^/person hour. For the aerial operation, we spent a total 168 person hours and recovered 126 of the estimated 256 carcasses (49%); a recovery rate of 0.04 carcasses/km^2^/person hour. Inclement weather forced us to conduct the aerial operation one day prior to planned without forewarning, and our full crew was not available. Therefore, during the first day with an insufficient carcass recovery crew (i.e., only two people per helicopter), we only recovered 23 of 153 (15%) of carcasses. During days two and three we had 5–6 people per helicopter and recovered 58 of 66 carcasses (88%) and 45 of 37 (121%) carcasses, respectively. We found more carcasses than expected on the third day because we underestimated piglets that were removed by the helicopter crew.

All ages of wild pigs were removed by each method, although the distributions of ages were dissimilar among the methods (Figure 6). For trapping, we mostly removed piglets (<6 months). For toxic baiting, we mostly removed juvenile and young adult wild pigs (∼6–24 months). For aerial operations we mostly removed piglets (<6 months) and old adults (>36 months).

**Figure 6.**
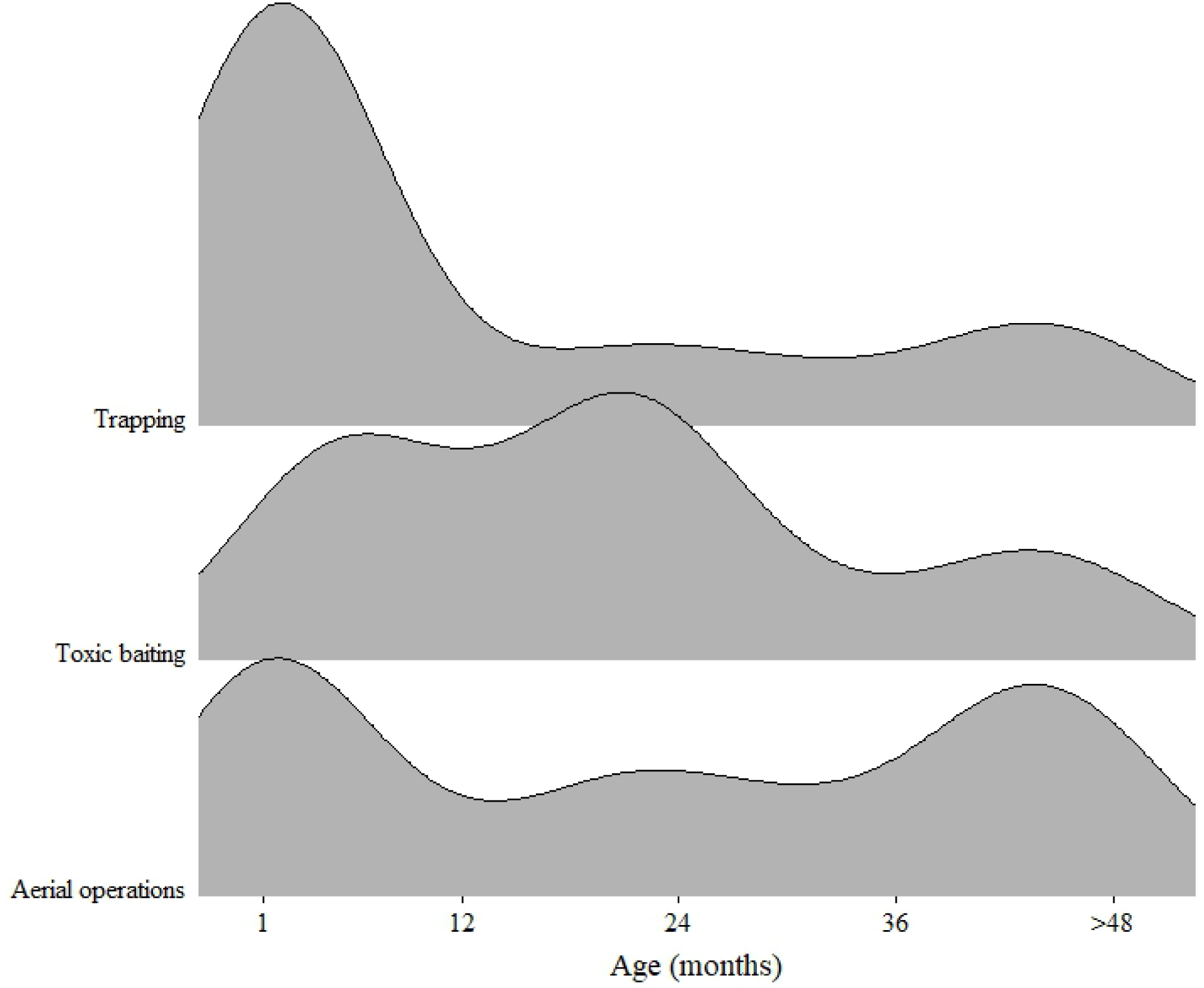
Distributions of estimated ages of wild pigs removed and carcasses recovered for each removal method for removing and recovering carcasses of wild pigs in northcentral Texas, USA during February–April 2023. We did not include ground shooting because too few animals were removed using this method.

## Discussion

There were strengths and weaknesses to all the removal methods for wild pigs, although aerial operations were the most time- and cost-efficient on a per wild pig basis. For quickly reducing the initial population density, aerial operations and trapping had similar cost efficiencies on a per 10% population reduction basis, although aerial operations achieved the reduction an average of 7.5 days faster (i.e., 177% faster than trapping). Toxic baiting appeared the be the least efficient method but was restricted by how much of the experimental bait we could deploy. We were required to cease toxic baiting with additional bait sites having been pre-baited and ready for toxic deployment, thus did not attempt to remove some wild pigs that we had invested into. Ultimately, we recognized that all methods could be used and intermixed during an FAD response, and our findings can help distinguish which methods might be prioritized for different situations and help establish budgets for emergency response activities. We discuss the operational strengths and weaknesses for each method below.

Aerial operations were most time- and cost-efficient, but also required the largest initial investment in equipment. However, as is the case for the US, some initial investments have already been made by the USDA/APHIS/Wildlife Services, thus any ongoing or future efforts could initially capitalize on existing equipment. Likely, additional investment would be required. The costs/wild pig that we observed ($146.97) were far more expensive than previous reports at $11.13/wild pig in Australia (Hone, 1990), and $23.90/wild pigs in the US (Bodenchuk, 2014), both converted to $USD and adjusted for inflation for the purposes of this manuscript. The added expense was likely related to carcass recovery in our effort and spending the extra time to intensively remove every wild pig possible, which wasn’t part of those previous studies. Regardless, costs of aerial operations can vary widely based on population density of wild pigs (Davis et al., 2018; Bengsen et al., 2022), and get exponentially more expensive as population densities are reduced (Fischer et al., 2020) or as the length of operations increases. The effectiveness of aerial operations relies on the flight crew being able to locate wild pigs, which our rangeland landscape during the late winter season prior to green-up was ideal. Other landscapes and seasons may be less efficient. We also conducted a quick, 3-day effort which was ideal for removing naive animals, but a FAD response would likely be conducted over months or even longer than a year (Pepin et al., 2022). Wild pigs may learn to avoid helicopters by hiding underneath cover (McCann et al., 2004). Despite this, wild pigs tended to not scatter outside of their normal space-use, at least for a short duration (Fischer et al., 2016), therefore incorporating another method of removal during an aerial operation could be valuable. We also found that aerial operations were also most efficient at quickly reducing the initial densities of wild pigs, which would be valuable to slowing/stopping disease transmission quickly, especially if followed-up or simultaneous with other methods. Additionally, we observed that removing small wild pigs (i.e., piglets) from the air was difficult, furthering the need for another removal method. Finally, we found that ≥3 carcass recovery crews were needed to keep up with the numbers of wild pigs that were being removed by a helicopter. We unexpectedly initiated the aerial operation one day early because of forecasted high winds later in the week, which left us with only one carcass recovery crew per helicopter during the first day. We only recovered 15% of the carcasses that first day, but 88–121% the subsequent days with additional crews.

Trapping was the second most time- and cost-effective method for removing wild pigs. This was also the most secure way of recovering carcasses. Our rates of capture/person hour (0.14) were less than previously reported rates (0.64-2.3; Gaskamp et al., 2021), likely because our trapping success declined as the population was reduced in our focal area. Our costs/wild pig ($244.32) were also greater than previous reports ($61.41; Bodenchuk, 2014), likely for the same reason. We also found that the costs to reduce comparable densities of wild pigs via trapping or aerial operations were similar, although trapping took much longer. We expect this similarity could be related to multiple factors, including: 1) we attempted whole sounder removal as much a possible which removed entire groups of pigs at once, 2) we learned to target where the most wild pigs were living by monitoring the area intensely with bait and trap sites, and 3) recovering carcasses took very few personnel hours with trapping. However, trapping was the most physically demanding removal effort, with 84% of person hours spent deploying bait, building, and tearing down traps. Also, we found that trapping was the most difficult on our personnel because working in the field all day followed by monitoring cellular traps at night was not sustainable, suggesting personnel dedicated to night-shift trap monitoring would be ideal. Additionally, trapping was the hardest on our vehicles because the traps were heavy and the terrain was rough, which resulted in frequent broken OHVs. There were also challenges with accessing traps following heavy rain (e.g., deep mud and road washouts), thus we had to anticipate which traps we could access before setting the traps. Similarly, some traps were inaccessible for multiple days and could not be rebaited, which increased acclimation time for wild pigs. Finally, trapping success declined with time, likely because the density of wild pigs was reduced, but also because some animals seemed educated against using traps after a negative experience (Snow et al., 2022). Another method of removal would be needed to overcome this.

The experimental sodium nitrite-toxic baiting was the most expensive on a per wild pig basis and was almost twice as expensive to achieve the same reduction in density as trapping or aerial operations, albeit the costs we reported are likely greater than would be realized under non-experimental conditions. To our knowledge, our costs/wild pigs are the first published for a SN-toxic bait and should be considered conservative costs. The EUP restrictions did not allow us to deploy the toxic bait in more than 11 locations, whereas we could have deployed more toxic bait with minimal effort and removed more wild pigs to increase the cost- and time-efficiency. We also could have deployed the toxic bait faster (e.g., ∼2–5 days faster) at some bait sites, but used a conservative approach considering we were obligated to give the experimental bait the best evaluation possible. A simulation study on Kangaroo Island, Australia estimated removing 0.31 wild pigs/person hour (Hamnett et al., 2023) using a similar SN-toxic bait, which was 3.4 times greater than the rate we realized. However, our rate included hours spent building cattle fences and conducing carcass recovery which accounted for ∼67% of our effort hours. Despite the costs, we identified some benefits of toxic baiting that make it a valuable tool for FAD response, including the equipment was smaller and lighter than trapping and fewer people were required to pre-bait and build the toxic bait sites (especially if cattle fencing was not required). Also, the absence of traps could render the wild pigs less wary. Conversely there were downsides to toxic baiting that should be considered. After the SN-toxic bait was deployed, locating carcasses required 2–4 people per toxic bait site to walk transects, and we only found ∼72% of the carcasses. Most recovered carcasses were ≤500 m from the bait sites, but we expect the carcasses we could not find were farther away. Indeed, previous studies found a small proportion of GPS collared wild pigs were found ≥1,000 m away from bait sites (Snow et al., 2024), which would be nearly impossible to find under normal circumstances. Further, the experimental SN-toxic bait had some risks for non-target animals that may not be acceptable in some situations (Snow et al., 2021a; Snow et al., 2021b). Finally, other toxic baits are available for wild pigs in different regions of the world. In the USA, a new toxic bait will be used operationally for the first time in 2024, containing the active ingredient warfarin, but deployment of this bait takes ∼10 weeks to remove wild pigs (Poché et al., 2018), therefore is likely to slow for an effective FAD response. Toxic baits containing sodium fluoroacetate (compound 1080) are used Australia and New Zealand (Twigg et al., 2005; Bengsen et al., 2017), but are losing support because of risks to non-targets (Green and Rohan, 2012; Warburton et al., 2022).

Ground shooting was the least effective method we tested for removing wild pigs. We did not attempt ground shooting as much as the other methods primarily because our study area was dense cedar and mesquite, and locating animals for ground shooting was not time efficient. More open landscapes may offer better success. Bodenchuk (2014) estimated that ground shooting cost $36.52/wild pig adjusted for inflation, but similarly did not remove many wild pigs using this method. Conversely, using a simulation model Hamnett et al. (2023) found ground shooting was a cost-effective approach, but estimated a much greater rate of removal (0.22 wild pigs/person hour) which was 5.5 times greater than we achieved. We suggest using ground shooting strategically, such as for trap-shy wild pigs, and not relying on it as major component of an FAD response.

Another substantial finding throughout this effort was that nearly 25% of our time was spent conducting logistical activities, which was greater than we anticipated. Despite this, the time we spent on logistical activities still seemed insufficient for fully addressing our needs. During a real FAD response, which would likely be larger in scope and time than our effort, anticipating and planning for logistical time would be critical.

We were surprised to observe differences in the age classes of wild pigs removed by each method, although we suspect the timing of a birth pulse likely confounded these results. We observed females farrowing during late February–early March which overlapped the timing of our removal methods, especially the toxic baiting (i.e., 03–07 March). During that time females had reduced movements, making them and their piglets difficult to remove (Snow et al., 2024). For trapping, our efforts extended from mid-March through April, thus the piglets had time to leave their farrowing nest and were likely naive, which increased our capture success of those younger animals. For aerial operations, this was the only method with increased number of older animals removed. We suspect aerial operations may be the most resistant method to learned-avoidance by wild pigs– where they learn to avoid removal activities because of previous negative experiences (e.g., Snow et al., 2022). We also suspect we battled learned avoidance in wild pigs with trapping and toxic baiting, because there was occasional trapping prior to our study by ranch employees that likely exposed some wild pigs to bait with negative experiences (e.g., Lewis et al., 2022; Kilgo et al., 2023). Finally, the helicopter crews reported difficulty in removing piglets despite our data indicating they removed a lot relative to other age classes. Our observations while on the ground suggested that aerial operations did miss some piglets, which exemplifies the importance of simultaneously trapping. Ultimately, we wished we had initiated our removal efforts ∼2 months earlier, before farrowing and a pulse of piglets were on the landscape.

There are other strategies for FAD responses with wild pigs that we did not evaluate but may be important to incorporate. Quickly erected fencing to prevent wild pigs from leaving an area while removal activities are ongoing would reduce further spread of an FAD and might increase effectiveness of removal activities (Lavelle et al., 2011). Fencing was implemeted to reduce the spread of ASFv in South Korea and found to be effective (Jo and Gortázar, 2021). Additionally, an oral vaccine for ASFv is under development for wild boar in Europe (Barasona et al., 2019; Sánchez et al., 2019), and strategies for delivery to the population are being established (e.g., Gervasi et al., 2024). Incorporating fences and vaccination programs into emergency responses may further reduce the risks of FADs spreading and should be evaluated for time- and cost-efficiency.

### FAD Response Implications

In a FAD response where recovery of wild pig carcasses is important, aerial operations and trapping could primarily be used together as the most time- and cost-efficient methods, with some caveats. For aerial operations, there needs to be enough carcass recovery crews to keep pace with helicopters, if high carcass recovery rates are desired or required to control an outbreak. In our case, ∼3 recovery crews (of two people each, each with one OHV) plus a coordinator (radio communication, plotting, and dispatching coordinates) per helicopter would have been minimally sufficient. In addition, trapping could be used to remove any wild pigs that become educated against the aerial operations, are too small to be effectively removed by aerial operations, or are conduct removals in areas where helicopters cannot safely or efficiently access. But, excessive trapping may generate some wary wild pigs that are difficult to remove. Toxic baiting should be considered if it is deemed more important to remove wild pigs than to recover the carcasses, such as when the population density becomes very low, and the animals are wary against the above methods. Ground shooting should be considered as the landcover or terrain allows or to strategically target wary wild pigs. Finally, we recommend having personnel dedicated to logistical tasks in the field, for staying organized and managing equipment during an entire FAD response. Establishing camera traps to support estimation of changes in density was valuable for comparing removal methods and would also be useful during an emergency response to determine if density reduction has been sufficient to reduce on-going transmission.

Ultimately, FAD emergency responses to diseases such as ASFv in wild pigs would likely cover a much larger area than our study. For example, Pepin et al. (2022) suggests intensive control of wild pigs in areas ≥300 km^2^ may be required, which is 5–15 times larger than any of our treatment areas. In addition, deployment of fencing to contain infected wild pigs, and/or deployment of oral vaccine baits would require substantial time and costs which we did not include here. Finally, we ceased control efforts as our resources for this study were limited, but a FAD response would likely attempt to reduce the population lower than our study accomplished. For these reasons, the magnitude of a true FAD response would likely be exponentially greater than reported here, and our results indicate that time and costs will be very high.

## Acknowledgments

The research was supported by NIFA-AFRI Agricultural Biosecurity program (GRANT13428587) and the United States Department of Agriculture. The findings and conclusions in this publication are those of the authors and should not be construed to represent any official US Government determination or policy. Mention of commercial products or companies does not represent an endorsement by the US government. We thank H. Ownbey and G. Studdard for access to private property. We thank Animal Control Technologies Australia for providing the placebo and SN-toxic bait. We thank multiple people from Texas Wildlife Services and Texas Parks and Wildlife for assisting with data collection. We thank anonymous reviewers for their comments on this manuscript.

