## Supplemental Table 1 for "Comparing efficiencies of population control methods for responding to foreign animal disease threats in wild pigs"

**Supplemental Table 1.** Estimated density and abundance (and 95% CIs) of wild pigs in pre-, during, and post-removals by method for wild pigs in northcentral Texas, USA during February–April 2023.

| DENSITY(km^2^) | Pre | During | Post |
| --- | --- | --- | --- |
| Trapping | 8.58 (5.62–13.1) | 3.09 (1.88–5.09) | 0.87 (0.37–2.02) |
| Toxic baiting | 4.51 (3.21–6.33) | NA | 3.46 (2.41–4.97) |
| Aerial operation | 12.96 (10.6–15.83) | NA | 6.25 (4.81–8.11) |
| Control | 1.55 (0.84–2.83) | 2.31 (1.53–3.49) | 0.92 (0.45–1.88) |
| ABUNDANCE |  |  |  |
| Trapping | 307 (201–469) | 111 (67–182) | 31 (13–72) |
| Toxic baiting | 247 (176–347) | NA | 190 (132–272) |
| Aerial operation | 489 (400–598) | NA | 236 (182–306) |
| Control | 77 (42–142) | 115 (76–175) | 46 (22–94) |
