## Supplemental Figure 1 for "Comparing efficiencies of population control methods for responding to foreign animal disease threats in wild pigs"

**Supplemental Figure 1.** Estimated number of animals removed (i.e. change in abundance) pre- and post-treatment compared with observed number of animals removed for wild pigs in northcentral Texas, USA during February–April 2023. Blue are mean estimates with 95% CIs obtained from time-to-event models and gray indicates known number of animals removed.

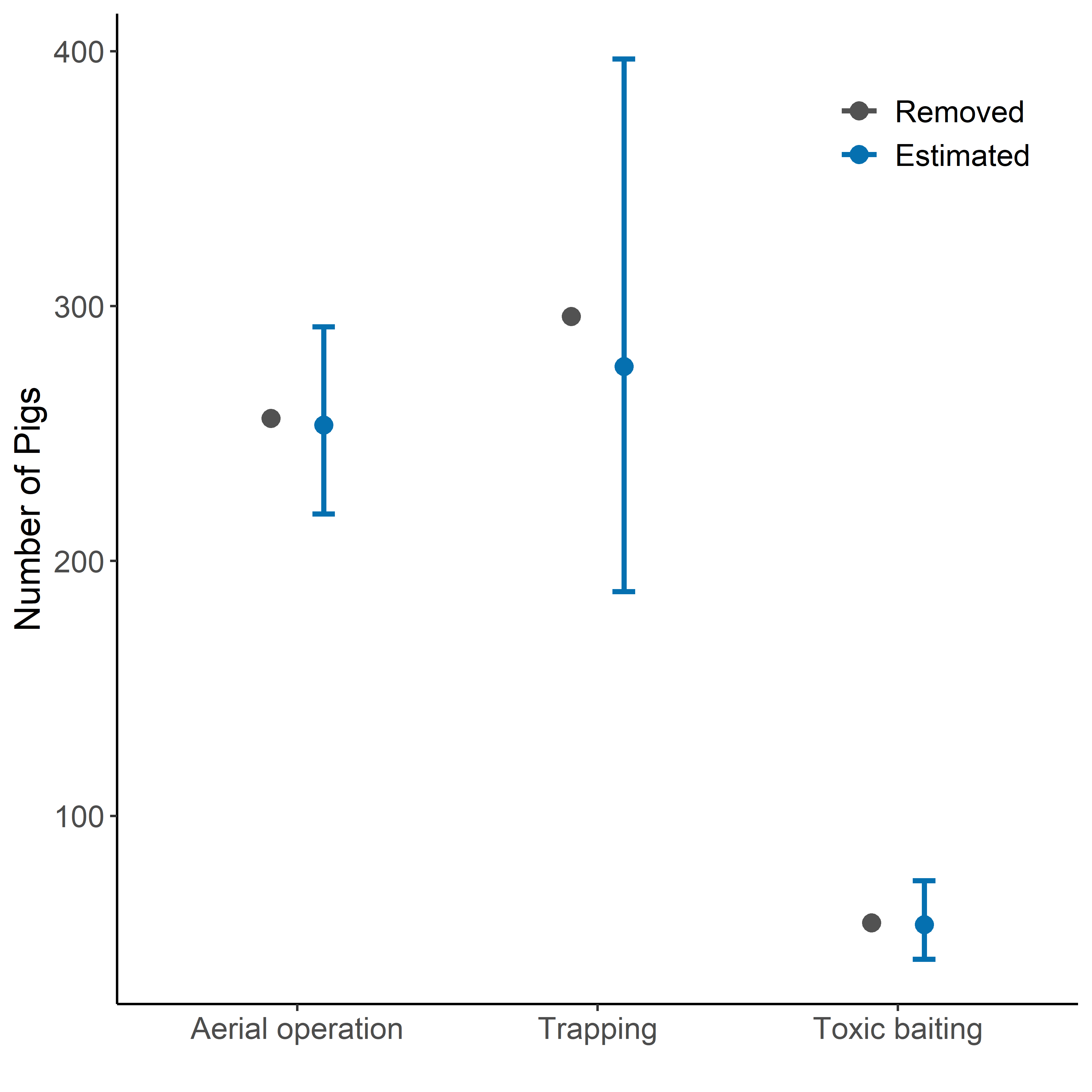
